# Computational Investigation of Increased Virulence and Pathogenesis of SARS-CoV-2 Lineage B.1.1.7

**DOI:** 10.1101/2021.01.25.428190

**Authors:** N. Arul Murugan, Prashanth S. Javali, Chitra Jeyaraj Pandian, Muhammad Akhtar Ali, Vaibhav Srivastava, Jeyakanthan Jeyaraman

## Abstract

New variants of SARS-CoV-2 are being reported worldwide. More specifically, the variants reported in South Africa (501Y.V2) and United Kingdom (B.1.1.7) were found to be more contagious than the wild type. There are also speculations that the variants might evade the host immune responses induced by currently available vaccines and develop resistance to drugs under consideration. The first step of viral infection in COVID-19, occurs through the interaction of receptor binding domain (RBD) of the spike protein with peptidase domain of the human ACE-2 (hACE-2) receptor. So, possibly the mutations in the RBD domain of spike protein in the new variants could modulate the protein-protein interaction with hACE-2 receptor leading to the increased virulence. In this study, we aim to get molecular level understanding into the mechanism behind the increased infection rate due to such mutations in these variants. We have computationally studied the interaction of the spike protein in both wild-type and B.1.1.7 variant with hACE-2 receptor using combined molecular dynamics and binding free energy calculations using molecular mechanics-Generalized Born surface area (MM-GBSA) approach. The binding free energies computed using configurations from minimization run and low temperature simulation show that mutant variant of spike protein has increased binding affinity for hACE-2 receptor (i.e. ΔΔG(N501Y,A570D) is in the range −20.4 to −21.4 kcal/mol)The residue-wise decomposition analysis and intermolecular hydrogen bond analysis evidenced that the N501Y mutation has increased interaction between RBD of spike protein with ACE-2 receptor. We have also carried out calculations using density functional theory and the results evidenced the increased interaction between three pairs of residues (TYR449 (spike)-ASP38 (ACE-2), TYR453-HIE34 and TYR501-LYS353) in the variant that could be attributed to its increased virulence. The free energies of wild-type and mutant variants of the spike protein computed from MM-GBSA approach suggests that latter variant is stable by about −10.4 kcal/mol when compared to wild type suggesting that it will be retained in the evolution due to increased stability. We demonstrate that with the use of the state-of-the art of computational approaches, we can in advance predict the more virulent nature of variants of SARS-CoV-2 and alert the world health-care system.

## 1 Introduction

A distinct phylogenetic cluster of Severe Acute Respiratory Syndrome Coronavirus-2 (SARS-CoV-2) referred as VUI 202012/01 (Variant Under Investigation, year 2020, month 12, variant 01) belonging to the lineage B.1.1.7 was recently identified in the United Kingdom through viral genome sequencing.^1^ It was found that this variant has unusual multiple mutations especially in the spike protein namely N501Y, A570D mutations in receptor binding domain (RBD) and H69/V70, 144/145 deletion mutations in S1 sub unit besides P681H, T716I, S982A, D1118H mutations in S2 sub unit.^2^ Driven by the increased infectiousness of this new strain, about 50,000 individuals per day had been infected in the United Kingdom till first week of January 2021 with the first case being identified on 20th September, 2020.^3^ VUI 202012/01 cluster varies by 29 nucleotide substitutions from the Wuhan strain that is much higher than the recent molecular clock predictions of about 2 substitutions per month.^1^ The non-synonymous mutations in the spike protein for VUI 202012/01 variant is higher than the expected random mutations and about 27 % of 22 nucleotide substitutions in the S-gene are acquired from common ancestor next strain clade 20B.^4^ Phylogenetic analysis of this variant evidenced 14 non-synonymous mutations involving amino acid alterations; 6 synonymous non-amino acid altering mutations and 3 deletion mutations.^1^ The six synonymous mutations include five mutations in ORF1ab namely C913T, C16176T, C14676T, C5986T, C15279T and one mutation in M gene viz. T26801C.^1^ Three deletion mutations in VUI 202012/01 variant comprise of i) several spontaneous double deletions (69/70 deletion) that possibly cause conformational change in the spike protein; ii) spontaneous multiple mutation (P681H) near high variability S1/S2 furin cleavage site and iii) open reading frame 8 (ORF8) stop codon (Q27stop) mutation.^5^ The N501Y mutation occurring in the RDB of spike protein is proposed to increase the binding affinity of the spike protein to murine and human angiotensin-converting enzyme-2 (ACE-2) receptors.^6,7^ Accumulation of high mutation rates within short time periods suggest that this variant might not have emerged from gradual accumulation of mutations but could have arisen by selection pressure through intrapatient virus genetic diversity after convalescent plasma treatment.^8^ The probable sources that has given rise to this variant with multiple mutations can be explained by three possible processes namely i) prolonged SARS-CoV-2 infection in immune-deficient/suppressed patients leading to accumulation of immune evading mutations at higher rate; ii) adaptation process of SARS-CoV-2 as in the case of Y453F mutation or 69/70 deletions that have occurred in different animal species followed by zoonotic transmission to human host and iii) antibody-mediated selective pressure creating multiple genetic changes in SARS-CoV-2 through direct selection or by genetic hitchhiking.^9,10^ Experimental studies have suggested that N501Y mutation in the RBD region of spike protein at position 501 can increase its affinity to the ACE-2 receptor.^11^ Animal studies using mouse model have also evidenced increased infectivity and virulence of N501Y mutation in SARS-CoV-2.^12^ Moreover, mutation in the residue P681H that creates furin cleavage among S1 and S2 sub-units of the spike protein resulted in enhanced SARS-CoV-2 variant transmission and entry into host respiratory epithelial cells in animal models.^13,14^ The double deletion mutation involving amino acids at sites 69-70 in N terminal domain of the spike protein is a recurrent mutation that has been recorded in multiple lineages linked to various RDB mutations.^15^ High frequency of the recurrent deletion mutations in RDB region of the spike protein were evidenced in Y453F mutation in Denmark’s mink-associated Y453F mutant outbreak and N439K mutation associated infections in human.^16^ Other significant mutation in B.1.1.7 variant occurs in ORF8 Q27 stop codon that truncate and inactivates ORF8 protein favouring further downstream mutations.^17^ For instance, the Singaporean strain had double deletions mutations leading to ablated ORF8 and truncated ORF7b expression besides 382nt deletion associated with reduced post-infection inflammation with milder clinical infection.^17^ Several studies have also reported that ORF8 deletion mutation has only modest effect on the replication of SARS-CoV-2 in the primary airway cells of human when compared to virus without this ORF8 deletion.^18^ Therefore, mutations in the critical residues of the RBD region plays a very important role in enhancing the interactions between the spike protein of SARS-CoV-2 and the ACE-2 receptor.^19^ Presence of F486 in RBD site instead of usual I472 can result in the formation of strong aromatic-aromatic interactions.^20^ This aromatic interaction is formed between Y83 of ACE2 and E484 in CTD, instead of ionic interactions between P470 in RBD and K31 which provide higher affinity for receptor binding. ^21^ Though there is no evidence that B.1.1.7 variant could produce severe illness compared to other SARS-CoV-2 strains, the following possible implications has been suggested for the new variant. The UK reports that the new variant has increased transmissible rate of about 70 % besides increase in reproductive number by 0.4. ^22^ The UK variant was also found to have impact on diagnosis of SARS-CoV-2 as the deletion mutations in 69-70 can cause negative result from S-gene in RT-PCR assay.^23–26^ Mutations in the RBD of spike protein or other surface structures can alter the antigenic property of new SARS-CoV-2 variant that could lead to reduction in neutralization activity of antibodies and thereby resulting in higher risk of reinfection or decreased effectiveness of vaccines. ^27^ Therefore, there is an urgent need to study the mutation pattern of SARS-CoV-2 and mechanism of virulence or pathogenesis of new variant B.1.1.7 to develop effective therapeutics. Based on the above considerations, in the present study, we have comprehensively investigated the multiple mutations in the genome of new SARS-CoV-2 variant B.1.1.7 besides its potential role in viral entry, replication or pathogenesis in comparison with the wild-type Wuhan strain and less contagious variants. This study also includes mutation pattern analysis, phylogenetic study and the possible sources of new variant B.1.1.7 through genomic analysis of SARS-CoV-2 sequences retrieved using Nextstrain data from GISAID for different geographical locations like India, Nigeria, United Kingdom and United Arab Emirates. Here, molecular dynamics simulation, free energy calculations and electronic structure theory based analysis of interactions of the residues from RBD domain of the spike protein and the ACE-2 receptor were performed to explore the stabilizing interactions in the formation of protein-protein complex and the effect of mutations (occurring in the RBD domain of the spike protein) in modulating complex stability. The findings of the study will enhance the understanding of genomic properties of new variant B.1.1.7 and the possible implications of its mutations on the viral protein structure that will offer a clue for futuristic development of novel therapeutics or potent vaccine against SARS-CoV-2.

## 2 Results and Discussion

The binding free energies based on MM-GBSA approach^28,29^ have been presented for configurations corresponding to minimization run and low temperature simulation (30 K and 1 atm pressure). The results corresponding to low temperature have been used since the entropic contributions are less significant in this condition. Further we will be able to compare the results from force-field methods to interaction energies computed using approaches based on quantum mechanics (QM). The binding free energies for the RBD domain of the spike protein with the ACE-2 receptor^30^ are given in Table 1. In particular, two cases namely wild type and mutant variants of the spike proteins were considered as receptors. As we see, the binding free energy for mutant (using the configuration from the minimization) are lower by about 20 kcal/mol when compared to wild type spike protein (will be referred as set-1). The trend remains the same even in the case of calculations using trajectories corresponding to low temperature simulation. Now the difference between the binding free energies for mutant when compared to wild type with ACE-2 receptor is about −21.0 kcal/mol (will be referred as set-2). Interestingly, the binding free energies obtained in this set of calculations are quantitatively lower when compared to results from minimization run. It has to be attributed that the low temperature simulation allows the system to sample configurational phase space which corresponds to minimum free energies. The binding free energies have the contributions from van der Waals and electrostatic interactions between spike protein-ACE-2 receptor and the polar and non-polar solvation free energy contributions. ^28,29^ The latter two terms account for the change in free energies associated with protein-protein complex formation in aqueous solvent like environment. The effect of solvent is to screen the charges on the amino acids and so the polar solvation free energy contributions generally are positive. ^31^ The electrostatic interactions and van der Waals interactions are contributing to the stabilization of the complex formation. The overall, electrostatic interactions are against the complex formation as the sum of electrostatic interaction and polar solvation free energies are positive. The values are respectively 102 and 77 kcal/mol for wild type and mutant variant of the spike protein with ACE-2 receptor (we refer to the results corresponding to set-2) and the complexes will be referred as wt-spike:ACE-2 and mt-spike:ACE-2 respectively. Therefore, the protein-protein complexation is driven by hydrophobic interactions between the residues in the spike protein with residues in the ACE-2 receptor. The van der Waals interactions are as much as −146 and −142 kcal/mol for the wild type and mutant variant of the spike protein with ACE-2 receptor. Interestingly, the N501Y mutation has lowered the van der Waals interaction (by about 4 kcal/mol while benefited due to increased electrostatic interaction by about 25 kcal/mol). The non-polar solvation contribution to the binding free energies are almost comparable in both cases even though its overall effect is to stabilize the complex formation.

**Table 1:** Various contributions to the binding free energies of wild-type and mutant variants of the spike protein with the ACE-2 receptor. The free energies were computed using MM-GBSA approach and the energies are in kcal/mol.

| System | $\Delta E_{vdw}$ | $\Delta E_{elec}$ | $\Delta G_{GB}$ | $\Delta G_{SA}$ | $\Delta G_{binding}$ |
| --- | --- | --- | --- | --- | --- |
| **Minimization** |  |  |  |  |  |
| wt-spike | -83.55 | -1042.27 | 1092.09 | -12.66 | -46.39 |
| B.1.1.7 spike | -91.70 | -979.95 | 1018.80 | -13.98 | -66.83 |
| $\Delta\Delta G(\text{N501Y,A570D})$ | | | | | **-20.44** |
| **Low temperature simulation** |  |  |  |  |  |
| wt-spike | -145.57 | -1118.60 | 1220.65 | -18.99 | -62.51 |
| B.1.1.7 spike | -142.18 | -998.37 | 1075.55 | -18.89 | -83.90 |
| $\Delta\Delta G(\text{N501Y,A570D})$ | | | | | **-21.39** |

We have analysed the residue-wise contributions to the binding free energies and this analysis has been carried out for both the spike protein and the ACE-2 receptor. Since our focus is to investigate the effect of mutations on the binding free energies in the complex formation, we have only presented the results for the spike protein. As we discussed earlier, we have only considered two mutations occurring in the RBD domain namely N501Y and A570D. The latter mutation occurs in the N-terminal domain which is located far away from the interfacial region involved in the protein-protein interaction (Refer to Figure 1a). So, it is expected that the latter mutation has no or less significant effect on the protein-protein interaction and the former one is supposed to have major contribution to difference in binding free energies between the wild-type and mutant variants of the spike protein with ACE-2 receptor. Figure 2a shows the residue wise contributions to the binding free energies from the residues of the spike protein and both wild type and mutant variants were considered. The numbering of the residues has been done as if they are in the whole spike protein even though we have studied only the RBD domain of this protein (322-580). In order to see the residues that are dominantly contributing to difference in binding free energies, we have computed △△E_Res_(_spike_)(wt → mt). One would expect major contribution from 501 residue as this corresponds to mutation site located in the interfacial region. However, the contributions from many other residues are not less significant which has to be attributed to the fact that the mutation occurring in 501 position also alters the distance between the other residues of the spike protein and ACE-2 receptor residues and so the interactions are altered significantly. How ever the major changes are observed for two residues 417 and 501. As expected the ΔΔE_Res(spike)_ (wt → mt) has stabilizing contribution (i.e. negative in magnitude and amounts to −7.7 kcal/mol) to the complex formation. It is also worth noticing that in the wild type variant, the 501 residue has destabilizing contribution to the complex formation (refer to the positive value for ΔE_Res(spike)_ for this residue in the case of wild type in Figure 2a). The contribution due to residue 417 is rather positive which suggests that this residue contributes to destabilization of complex formation in the case of N501Y mutant. In addition to stabilizing contributions from residue 501, there are other residues such as 486, 489, 493, 496, 500, 502 are stabilizing the mutant N501Y with a contribution < −1 kcal/mol.

**Figure 1:**
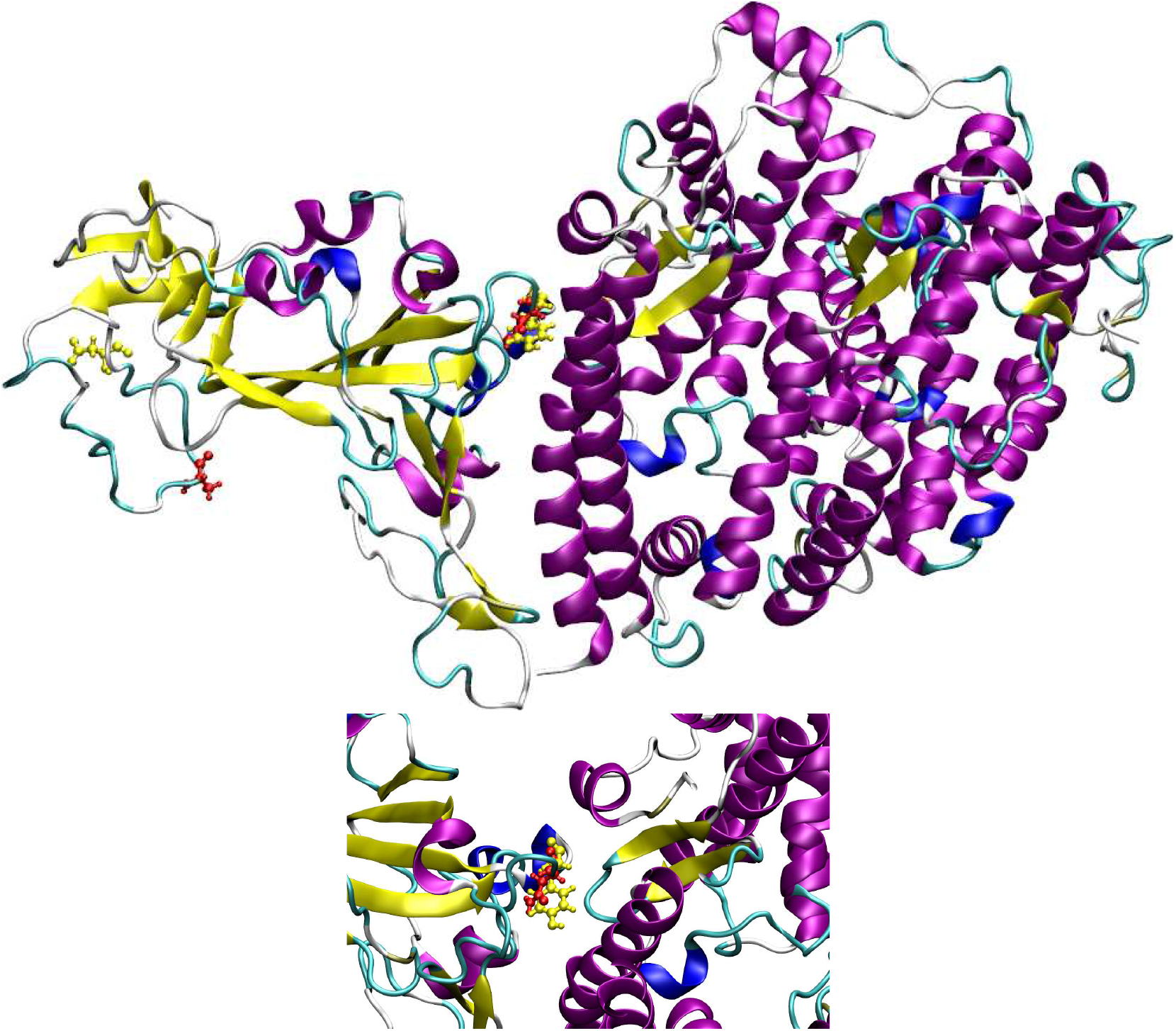
Three dimensional structure the spike protein:hACE-2 complex. The residues that are mutated in B.1.1.7 variant are shown in yellow color.

**Figure 2:**
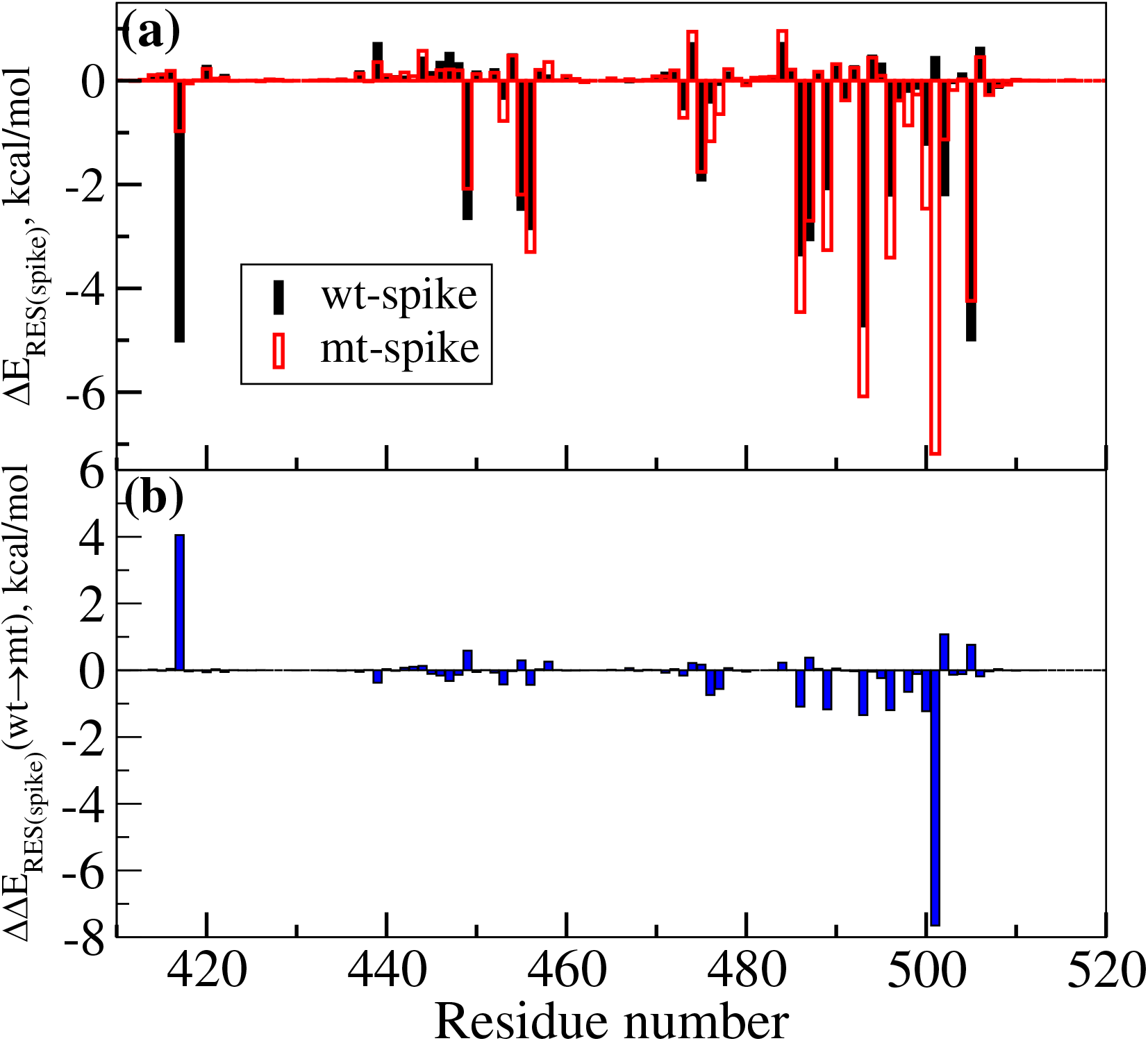
The spike protein residue wise contributions to the binding free energies with ACE-2 receptor. Results are shown for both wild type and B.1.1.7 variants of SARS-CoV-2 in *(a)* and the differences between these two values are shown in *(b)*

Further to address the stability of the two protein-protein complexes (i.e wt-spike:ACE-2 and mt-spike:ACE-2), root mean square displacement (RMSD), root mean square fluctuation (RMFS) and hydrogen bond analysis were carried out using the trajectories corresponding to 50 ns production run. The results for RMSD are given in Figure 3a and 3c while the RMSF results are given in 3b and 3d. The first configuration in the production run has been used as the reference for computing RMSD and the values are presented for the spike protein and the ACE-2 protein separately. The RMSD values are shown to fluctuate over a larger range for the spike protein when compared to ACE-2 receptor which indicates the former biomolecule undergoes larger conformational flexibility than the latter. It is expected behaviour as the RBD domain of the spike protein only has been studied in this work and the conformational stability (or structural stability) is reduced as we do not include the whole spike protein in trimeric form. ^32^ Another interesting observation is that both the spike protein and the ACE-2 receptor in the case of mutant variant are associated with the larger RMSD values suggesting that conformational fluctuations are increased due to mutation N501Y. We have seen that this mutation is associated with lowering of the binding free energies but it is interesting to see it also contributes to larger conformational flexibility. The increased conformational flexibility has been attributed to the increased ability for the spike protein to escape from the neutralizing potential of antibodies. ^33^ RMSF values shown for the spike protein suggest that the such conformational flexibility occurs rather over a small range of residues namely 527-580. In the case of the ACE-2 protein bound to mutant variant of the spike protein larger RMSF values were observed signifying that the mutation occurring in spike protein can induce changes in structure and dynamics of ACE-2 protein.

**Figure 3:**
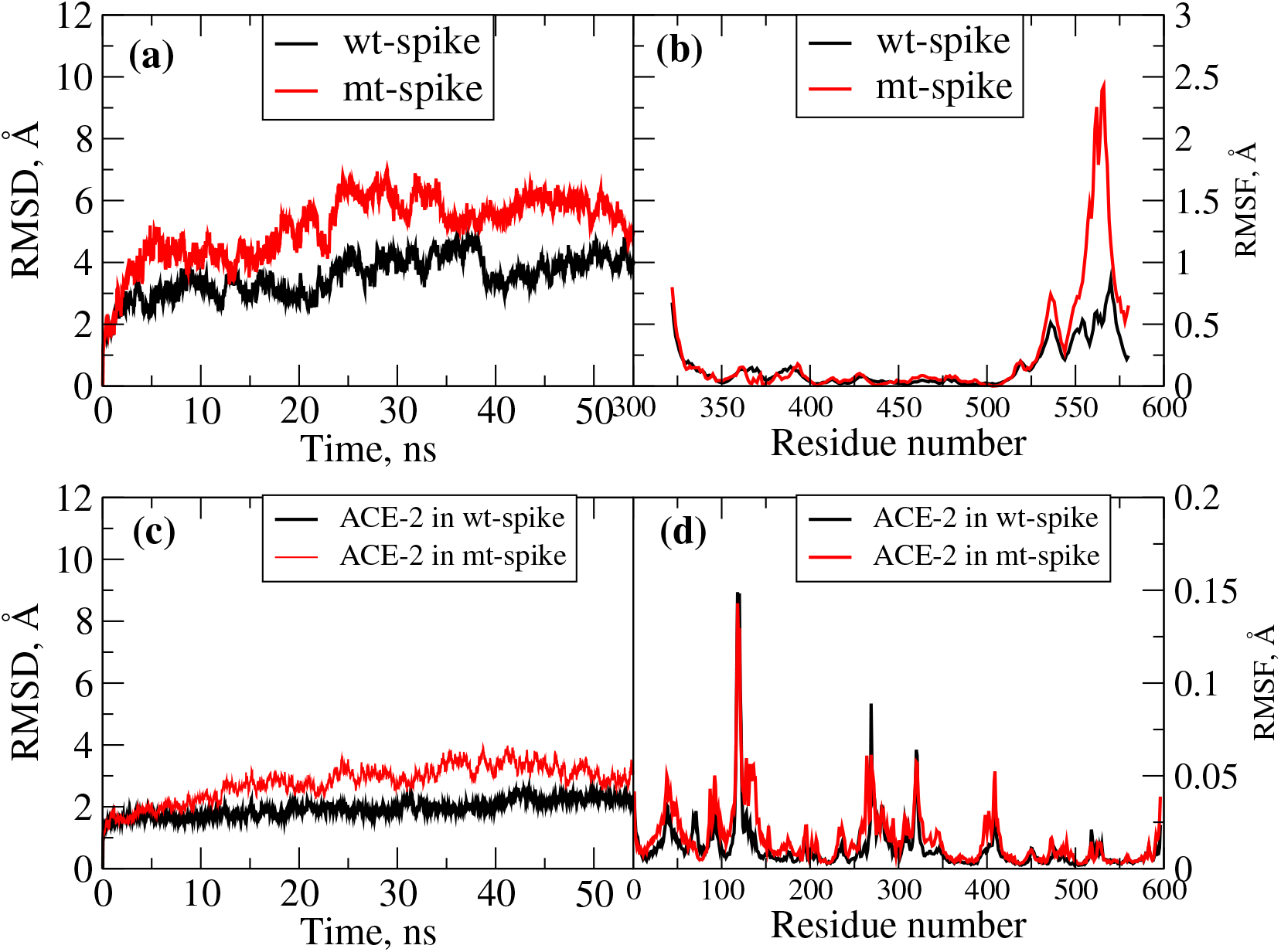
*(a)* RMSD as a function of time computed for wild type and mutant variant of the RBD domain of spike protein when it is bound to the ACE-2 receptor. *(b)* RMSF for wild type and mutant variant of the RBD domain of spike protein when it is bound to the ACE-2 receptor. *(c)* RMSD as a function of time computed for the ACE-2 receptor when it is bound to wt-spike and mt-spike proteins. *(d)* RMSF for ACE-2 receptor when it is bound to wt-spike and mt-spike proteins.

We have also analysed the intermolecular hydrogen bonds in the spike protein:ACE-2 complex for the two variants namely wild type and B.1.1.7 variant. The results are presented in Table 2. The residues involved in hydrogen bonds (hbonds), fraction of configurations the hbonds survive and average hydrogen bond geometries are provided in the Table. We have only listed H bonds that have survival percentage greater than 40%. This means these hbonds were observed at least in 40% configurations in whole trajectories. The hbonds having larger survival time are the most stable. As can be observed from Table 2, the number of hbonds with survival fraction of 0.4 are 6 in the case of wt-spike:ACE-2 complex. In the case of mutant variant the number of such hbonds increased to 8 which explains why the protein-protein complex is more stable in this case. It is interesting to note that the most stable hydrogen bond (with larger survival fraction) is not the same in the case of both wt-spike:ACE-2 and mt-spike:ACE-2 complexes. In the case of wt-spike:ACE-2 complex, the most stable hydrogen bond is formed between the TYR449 of the spike protein and ASP38 of ACE-2 receptor while for mt-spike:ACE-2 complex, it is established between ASN487 (of spike protein) and TYR83 (of ACE-2 receptor). Even though the residue in mutation site is not involved in hydrogen bonding, the residues in the vicinity of the mutation site such as THR500, GLY502, GLN498 are involved in hydrogen bonding. This explains why the ΔΔE_Res_ (wt – spike – mspike) were non-zero for the residues located in the vicinity of mutation site. As can be seen the hydrogen bonding formed by these residues in the wt-spike protein with ACE-2 are significantly affected by the mutation. There were two hydrogen bonds formed by THR500 in wild type spike protein which is reduced to single hydrogen bond in the case of mutant variant. Further the hydrogen bond formed by GLN498 in wild type is lost in the case of mutant. In the latter case, new hbonds are formed. Single mutation occurring at 501 site significantly alters the intermolecular hydrogen bonding pattern which stabilizes the protein-protein complex formation for N501Y mutant. Further, the N501Y leads to increased number of hbonds having larger survival time and this has to be attributed to the increased interaction between the spike protein and ACE-2 receptor.

**Table 2:** Hydrogen bond analysis carried out for the spike protein:ACE-2 complex. Only those hbonds with survival fraction greater than 0.4 are shown here. Also the avergage hydrogen bond geometries are shown.

| No | Acceptor | Donor | Fraction | $R_{H-bond}$ | $\theta_{H-bond}$ |
| --- | --- | --- | --- | --- | --- |
| **wt-spike:ACE-2 receptor complex** |  |  |  |  |  |
| 1 | ASP38 (ACE-2) | TYR449 (spike) | 0.9378 | 2.6814 | 164.2380 |
| 2 | HIE34 (ACE-2) | TYR453 (spike) | 0.7929 | 2.8201 | 162.9223 |
| 3 | ASP38 (ACE-2) | GLN498 (spike) | 0.7187 | 2.8274 | 164.3970 |
| 4 | LYS353 (ACE-2) | GLY502 (spike) | 0.6827 | 2.8685 | 161.4827 |
| 5 | ASP355 (ACE-2) | THR500 (spike) | 0.4387 | 2.6902 | 163.6439 |
| 6 | TYR41 (ACE-2) | THR500 (spike) | 0.4089 | 2.7926 | 161.8899 |
| **mt-spike:ACE-2 receptor complex** |  |  |  |  |  |
| 1 | ASN487 (spike) | TYR83 (ACE-2) | 0.9338 | 2.7036 | 159.1010 |
| 2 | TYR83 (ACE-2) | TYR489 (spike) | 0.8733 | 2.7876 | 159.7186 |
| 3 | HIE34 (ACE-2) | TYR453 (spike) | 0.7182 | 2.8324 | 162.9428 |
| 4 | LYS353 (ACE-2) | GLY502 (spike) | 0.5964 | 2.8780 | 162.8891 |
| 5 | TYR41 (ACE-2) | THR500 (spike) | 0.5867 | 2.8052 | 159.6009 |
| 6 | GLN24 (ACE-2) | ASN487 (spike) | 0.5507 | 2.8531 | 155.7901 |
| 7 | GLU35 (ACE-2) | GLN493 (spike) | 0.4276 | 2.8268 | 163.8225 |
| 8 | ALA475 (spike) | SER19 (ACE-2) | 0.4142 | 2.7294 | 160.5896 |

Finally we have analysed the interaction energies of the key residues using electronic structure theory to further validate the results from force-field based approaches. We have identified the pair of residues from the spike protein and the ACE-2 receptor appearing within a cut-off distance of 8 Å. The pair of residues separated by distances larger than this may not contribute to the stabilization of the protein-protein complex. So, they have not been considered for this analysis. In the case of configuration corresponding to minimization runs there were about 53 such pairs and the interaction energies between them and the total sum were obtained using two different approaches. In the first method, the energies of the three systems namely residues from spike protein, ACE-2 receptor and their complexes were obtained in aqueous solvent environment and the difference in energies of the complex and subsystems (i.e. spike protein and ACE-2 receptor centered residues) gives the interaction energies of the pairs of residues of protein-protein complex in aqueous environment. The interaction energies between the pairs of residues located on the spike protein and ACE-2 receptor were obtained from the expression as shown in equation 1. In particular, we employed M06-2X/6-31+G* level of theory^34^ and for solvent description SMD solvent model^35^ has been adopted. The interaction energies of the protein-protein complex were computed as sum over the interaction energies between the pairs of residues and the values are presented in Table 3. The sum over interaction energies for wt-spike:ACE-2 complex is −67.5 kcal/mol while for the mt-spike:ACE-2 complex is −86.4. The ΔΔE_spike:ACE–2_(wt – spike → mt – spike) corresponding to N501Y mutation is about −19.0 kcal/mol which explains why the B.1.1.7 mutation is more contagious when compared to wild type. In addition to the energetics of the subsystems and complexes in aqueous environment, this set of calculations also provide the solvation energies of these systems. These energies refer to the change in energies corresponding to the change of the environment from vacuum (or gas-like) to aqueous condition. These solvation free energies are referred as ΔGSolv_aqueous_ and the difference in solvation free energies of the complex to the subsystem solvation free energies are given in the Table 3.

**Table 3:** Interaction energies (ΔE_aqeuous_) computed by different approaches using electronic structure theory. The solvation energies (ΔSolv_aqeuous_) associated with complex formation has been computed as the difference in solvation free energies for the complex and for the subsystems.

| System | $\Delta E_{\text{aqueous}}$ | $\Delta \text{Solv}_{\text{aqueous}}$ | $\Delta E_{\text{BSSE}(\text{raw})}$ | $\Delta E_{\text{BSSE}(\text{cor})}$ | $\Delta E_{\text{1}_{\text{aqueous}}}$ |
| --- | --- | --- | --- | --- | --- |
| wt-spike | -67.51 | 191.71 | -259.09 | -243.69 | -51.98 |
| B.1.1.7 spike | -86.51 | 236.72 | -323.22 | -304.29 | -67.57 |
| $\Delta \Delta E(\text{N501Y}, \text{A570D})$ | **-19.00** | 45.01 | -64.13 | -60.60 | **-15.60** |

In the second method, the interaction energies were obtained using counterpoise correction method by employing M06-2X/6-31+G* level of theory. The values obtained using this method are presented in Table 3 as ΔE_BSSE(raw)_ and ΔE_BSSE(cor)_. In particular, the former one gives the ΔE as the mere energy difference between the complex and subsystems. However, the latter case the energies are corrected for basis sets superposition error ^36,37^ using counterpoise correction method. ^38^ Moreover, M06-2X level of theory has been often used to estimate the interaction energies in even weak intermolecular complexes where the major contributions are due to van der Waals type interaction. ^39^ Even density functional theory with dispersion correction is known to perform reasonably well to describe such interactions. ^39^ However, we have used M06-2X level of theory to describe the interactions between the pairs of residues in the interfacial region of protein-protein complex. All the electronic structure theory calculations were carried out using Gaussian09 software.^40^

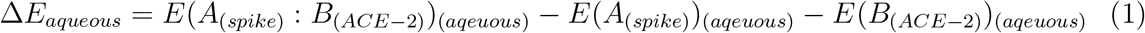

Since this approach, considers vacuum like environment to estimate the interactions between the residues, it tends to overestimate when the residues are oppositely charged or when a single residue is charged while the other residue is neutral. Using this approach, the sum over interaction energies computed for wt-spike:ACE-2 complex is −243.7 kcal/mol while for the mt-spike:ACE-2 complex is −304.3 kcal/mol (we are referring to BSSE corrected values). The ΔΔE_spike:ACE-2_ corresponding to N501Y mutation using the counterpoise correction method is about −60.6 kcal/mol which also explains why the B.1.1.7 mutation is more contagious when compared to wild type. As we mentioned this approach underestimates the interaction energies as the solvation effect is not accounted for. However, one can add to this the solvation free energies (i.e. ΔGSolv_aqueous_) which are computed in the first step and the sum has been presented as ΔΔE1_aqueous_ in Table 3. The change in interaction energy for the N501Y mutation is about −15.6 kcal/mol which is closer in magnitude to the values presented in the column ΔΔE_aqueous_. Two approaches namely the interaction energies obtained using the expression 1 and the interaction energies computed from the counterpoise method added together with solvation free energies (ΔGSolv_aqueous_) provide comparable results for the binding free energies for spike protein:ACE-2 receptor. Further both method shows that due to N501Y mutation the binding free energies are lowered by −19 and −15.6 kcal/mol. These results are consistent with the observation that B.1.1.7 mutant is 70% more contagious which has to be attributed to the increased interaction between the spike protein: ACE-2 receptor which is the first step in the infection.

We have also analysed interactions between the pairs of residues that majorly contribute to the total interaction energies. The pairs that contribute with interaction energies <-4.0 kcal/mol in the case of wt-spike:ACE-2 receptor are LYS417-ASP30, TYR505-GLU37 and TYR505-LYS353 and the ΔE_(spike):(ACE-2)_ for these pairs of residues are respectively, 6.8, −6.4 and −4.7 kcal/mol. In the case of mt-spike:ACE-2 complex, interestingly there are six such pairs: LYS417-ASP30, TYR449-ASP38, TYR453-HIE34, TYR501-LYS353, TYR505-GLU37 and TYR505-LYS353. Further the contributions to the total interaction energies from these residues are respectively, −6.3, −7.9, −4.5, −5.2, −6.3 and −4.7 kcal/mol. As can be seen the increased affinity for ACE-2 receptor in the case of mutant variant of spike protein is caused by the pairs, TYR449-ASP38, TYR453-HIE34 and TYR501-LYS353. Even though the mutation occurs in site 501 only, the effect is also propagated to other residues and this is an very interesting observation in this study.

We have also analysed whether the increased stability of the mutant may be a cause for the retainment and propagation of this specific variant. The free energy of the spike protein also will be able to can shed light into this aspect. So, we have also calculated the free energy of the both wt-spike and mt-spike proteins and listed in Table 4. Both total intramolecular energy (total energy of the protein due to bonded and non-bonded interactions between its residues) and solvation energies (both polar and non-polar contributions) are also shown. As can be evidenced the results corresponding to minimization run, the stability of mutant variant is increased by about 4.7 kcal/mol (i.e. the free energy of the mutant is lowered by this much value). The results from low temperature run also shows the same trend and suggest that mutant variant is more stable when compared to wild type by about −10.4 kcal/mol. The study also demonstrates one should be able to estimate the relative stability of spike proteins of different mutants to analyse whether it will be retained as the virus evolves.

**Table 4:** Various contributions to the free energies of wild and mutant variants of the spike protein in aqueous solvent environment. The free energies were computed using MM-GBSA approach and the energies are in kcal/mol.

| System | $E_{intra}$ | $G_{polar}$ | $G_{non-polar}$ | $G_{spike}$ |
| --- | --- | --- | --- | --- |
| **Minimization** |  |  |  |  |
| wt-spike | -6552.25 | -2394.82 | 105.24 | -8841.83 |
| B.1.1.7 spike | -6527.80 | -2427.62 | 108.89 | -8846.53 |
| $\Delta G_{spike}(N501Y, A570D)$ | | | | **-4.70** |
| **Low temperature simulation** |  |  |  |  |
| wt-spike | -6249.64 | -2362.61 | 94.04 | -8518.21 |
| B.1.1.7 spike | -6263.04 | -2361.27 | 95.70 | -8528.61 |
| $\Delta G_{spike}(N501Y, A570D)$ | | | | **-10.40** |

## 3 Computational Methods

### 3.1 Mutants preparation

The B.1.1.7 is reported to have as many as 23 mutations in the viral genome. However, only 8 of them occurred in the spike protein.^1–3^ Among these two are deletions occurring at positions HV 69-70, Y144 and others are N501Y, A570D, P681H, T716I, S982A and D1118H. As can be seen only one mutation is occurring in the RBD domain (N501Y) and another mutation is in the linker region between the RBD domain and NTD domain (A570D). Since the RBD domain of the spike protein is responsible for binding to the ACE-2 receptor, we have only studied the effect of these two mutations on the spike protein binding to the ACE-2 receptor. We have used PyMOL software^41^ to propose the structure for the mutant variant of the spike protein RBD domain.

### 3.2 Molecular Dynamics Simulation and MM-GBSA free energy calculations

Two systems of protein-protein complexes were considered in this study. (i) RBD domain of wild spike (will be referred as wt-spike) protein (322-580) with the ACE-2 receptor (19-614) (ii) double muted variant of spike protein (will be referred as mt-spike) with the ACE-2 receptor. As we mentioned above, the mutant spike protein differ from wild type in terms of the two mutations N501Y and A570D. The positions of the two mutations are shown in Figure 1. As can be seen the former mutation occurs in the RBD domain which is in direct contact with residues in the ACE-2 receptor (key residues are TYR41, GLY352, LYS353, GLY354, ASP355). The second mutation occurs in the NTD domain of the spike protein and does not have any role in dictating the binding affinity towards ACE-2 receptor. The three dimensional structure for the wt-spike:ACE-2 complex is based on the homology models build using the template structure reported for the complex in the PDB ID 7A91.^42^ The structure for the complex was build using SWISSMODEL server. ^43^ There are structures with better resolution reported for the ACE-2 receptor however, we chose this as template file as we wanted to propose the structure for the spike protein with residues in the range 322-580. The structure obtained from homology modeling has been used for preparing the input files for carrying out molecular dynamics simulations. The complex has been solvated with sufficient number of water molecules and neutralized with counter ions. The solvated complex structure for the wt-spike:ACE-2 complex has been energy minimized and followed by this low temperature simulations (at 30 K and 1 atm pressure) were carried out. Finally, MD simulations in ambient conditions (300 K and 1atm) were carried out. A similar protocol has been followed for mt-spike:ACE-2 complex except that before processing the PDB structure for the MD input file preparation, the two aforementioned mutations were introduced in the spike protein using Mutagenesis module of PyMOL software. All the finite temperature simulations were carried out isothermal-isobaric ensemble by using AMBER16 software.^44^ The time step for solving Newton’s equation of motion was set to 2fs. The time scale for the production runs for the wt-spike:ACE-2 and mt-spike:ACE-2 complexes was set to 50 ns. This set of simulations were carried out to study the stability of the complexes during a long time scale. In addition, the trajectories were used for computing various properties like RMSD, RMSF and hydrogen bond analysis. In particular, we were only analysed the intermolecular hydrogen bonds between the spike protein and ACE-2 receptor. The low temperature simulation and the structure corresponding to minimization run were used for computing the binding free energies using molecular mechanics-Generalized Born surface area approach (MM-GBSA). Even for the trajectory corresponding to finite temperature simulation at 300 K and 1 atm pressure the binding free energies were computed and the trends in them reflected the results corresponding to above two sets (even though quantitatively the values were in the higher side). The reason for using the configurations from the low temperature simulation is to avoid the need for computing the entropic contributions and in addition it allows us to directly compare to the energetics obtained from subsequent QM calculations. For the QM calculations, only those pairs of residues located in the interfacial region and are separated by a distance of 8 A were considered. Further, the residues were capped with hydrogens to meet with the valency requirements for carbon and nitrogen atoms after cutting along the peptide bonds.

## 4 Conclusions

Currently, a major attention is given to identify the location and spread of highly contagious variants of SARS-CoV-2 as it is feared that certain variants can escape the vaccination or develop resistance to drugs^45,46^ based therapy. Along this line the variant reported in South Africa and in United Kingdom (referred as B.1.1.7 variant) are reported to be more contagious and are reported to spread at 70% higher rate than the wild variant. It is not clearly established why these variants are highly contagious and we have undertaken a computational investigation to identify the mechanism behind the increased virulence and pathogenesis. The infection occurs through the first step that the spike protein binds to the ACE-2 receptor in human cell. So, naturally the mutations occurring in the RBD domain should give us details about the increased virulence of these variants. Therefore, we have studied the interaction between the wt-spike with ACE-2 receptor and mt-spike interaction with the same receptor using computational approaches. The calculated binding free energies clearly showed that the interaction between the mt-spike to ACE-2 receptor was increased by about −20 to −21 kcal/mol. We have also computed the change in interaction energies due to N501Y mutation using electronic structure theory based calculations and it is consistently found that the binding free energies for ACE-2 receptor are lowered by < −17 to 19 kcal/mol for the mutant variant of spike protein. The study clearly explains why the variants showed increased virulence and we attribute this to the more stable complex formation between the spike protein:ACE-2 receptor. We have also found the increased interaction between the certain pairs of residues (such as TYR449-ASP38, TYR453-HIE34 and TYR501-LYS353) and increased number of intermolecular hydrogen bonds to be responsible for this. We hereby, demonstrate that the computational modeling can provide valuable insight on how the mutations can modulate the protein-protein interaction in the spike protein:ACE-2 complex. Finally, we have also shown the mutant variant of the spike protein is stable by −4. to −10 kcal/mol when compared to wild type suggesting that it may be retained in the virus evolution.

## 5 Acknowledgments

This work was supported by the grants from the Swedish Infrastructure Committee (SNIC) for the projects “In-silico Diagnostic Probes Design” (snic2020-5-2). JJ gratefully acknowledges the MHRD-RUSA 2.0 [F. 24/51/2014-U, Policy (TNMulti-Gen), Dept. of Edn. Govt. of India] for the infrastructure facilities provided to the Department of Bioinformatics, Alagappa University. JJ thanks DST INDO-TAIWAN (GITA/DST/TWN/P-86/2019 dated: 04/03/2020) for the funding.

## 6 Author contributions statement

NAM and JJ designed the project. MD, binding free energy and QM calculations and analysis were carried out by NAM. Genomics analysis was carried out by MAA, VS, PSJ, CJP and JJ. Manuscript was written by NAM, CJP and JJ with contributions from all authors. All authors participated in the scientific discussion.

## 7 Supporting information

The supporting information includes details about genomics analysis and various mutations in B.1.1.7 variant when compared to wild type SARS-CoV-2. Further the sequences of wild type and sequence information for different mutants are provided.

